## Supplemental Materials for "Collective multicellular patterns arising from cadherin-linked cytoskeletal domains"

Note: equations and figures in the main text are referred to using the letter M preceding the equation or figure number, e.g Eq. (M4) and Fig. M4.

### I. MATHEMATICAL MODEL

#### A. Background of passively crosslinked actin

Suppose that the interior of each cell is comprised of 1) a background population of actin that is merely passively crosslinked having density $\rho_0$ and 2) a population of actin that is both actively crosslinked with motors and passively crosslinked having density $\rho$. The two populations of actin travel with the same velocity, as they are passively crosslinked. The internal stress is $\boldsymbol{\sigma}$ as given by Eq. (M3) in the main text and reiterated here:

$$\boldsymbol{\sigma} = \eta(\rho+\rho_0)^2\left(\nabla\mathbf{u} + (\nabla\mathbf{u})^T\right) - s_1(\rho+\rho_0)^2\mathbb{1} + s\rho\mathbb{1} \quad . \tag{1}$$

Meanwhile, the two populations of actin, densities $\rho$ and $\rho_0$ are in interchange with one another with rates $k_{\text{bind}}$ and $k_{\text{unbind}}$:

$$\frac{\partial\rho}{\partial t} + \nabla\cdot(\mathbf{u}\rho) = D\nabla^2\rho + k_{\text{bind}}\rho_0 - k_{\text{unbind}}\rho \tag{2}$$

$$\frac{\partial\rho_0}{\partial t} + \nabla\cdot(\mathbf{u}\rho_0) = D_0\nabla^2\rho_0 - k_{\text{bind}}\rho_0 + k_{\text{unbind}}\rho + V - B\rho_0 \quad . \tag{3}$$

Here $k_{\text{bind}}$ represents the rate at which motors bind to the background pool of passively crosslinked actin, converting $\rho_0$ to $\rho$, and $k_{\text{unbind}}$ represents the rate at which motors unbind from actively crosslinked actin, converting $\rho$ to $\rho_0$. The diffusion of coefficients of the actin species are $D$ and $D_0$. We assume that background actin experiences interchange with a large pool of actin monomers, with nucleation rate $V$ and degradation rate $B$. If we assume that $V$ and $B$ are fast compared to other processes in the system such as $k_{\text{bind}}$, $k_{\text{unbind}}$, diffusion, and velocities, then $\rho_0$ can be approximated by a constant $\rho_0 \approx V/B$, and Eq. (3) can be eliminated. The physical interpretation of $\rho_0$ would be that it is a roughly constant population of passively crosslinked filamentous actin that does not exert active forces, yet it experiences steric repulsion in the same way as does actively crosslinked actin. This simplification leaves only Eq. (2) to evolve $\rho$ as:

$$\frac{\partial\rho}{\partial t} + \nabla\cdot(\mathbf{u}\rho) = D\nabla^2\rho + \nu - b\rho \tag{4}$$

with $\nu \equiv k_{\text{bind}}\rho_0 \approx k_{\text{bind}}V/B$ and $b \equiv k_{\text{unbind}}$, presented as Eq. (M**??**) of the main text.

### B. Non-dimensionalization

We reiterate the equations of motion from the main text (Eqns. (M1)-(M2), (M6), and (M9)) here for quick reference.

$$\frac{\partial \rho}{\partial t} = D\nabla^2\rho - \nabla\cdot(\mathbf{u}\rho) + \nu - b\rho \tag{5}$$

$$\mathbf{0} = \nabla\cdot\Big(\eta(\rho+\rho_0)^2\Big(\nabla\mathbf{u} + (\nabla\mathbf{u})^T\Big) - s_1(\rho+\rho_0)^2\mathbb{1} + s\rho\mathbb{1}\Big) - \gamma(\rho+\rho_0)\mathbf{u} \tag{6}$$

$$\frac{\partial \psi}{\partial t} = -\frac{\partial}{\partial s^+}\Big(-D_\mathrm{e}\frac{\partial \psi}{\partial s^+} + v^+\psi\Big) - \frac{\partial}{\partial s^-}\Big(-D_\mathrm{e}\frac{\partial \psi}{\partial s^-} + v^-\psi\Big) + a\bar{\delta}(|s^+ - s^-|) - \lambda_0\psi \tag{7}$$

$$v^\pm = \frac{k(s^\mp - s^\pm)}{\mu_0(\rho^\pm + \rho_0)} + u^\pm \quad . \tag{8}$$

Here $\bar{\delta}(|s^+ - s^-|)$ represents the expression multiplied by $a$ in Eq. (M7), and we have substituted expressions for $k_{\mathrm{on}}$ and $k_{\mathrm{off}}$ into Eq. (7); as the piecewise limits at $\ell_\mathrm{M}$ are disregarded here for notational simplicity, Eq. (7) applies only to $|s^+ - s^-| \leq \ell_\mathrm{M}$. There is also a boundary condition that is non-trivial to find in non-dimensional form; this is a combination of Eq. (M4) and Eq. (M10):

$$\hat{\mathbf{t}}\cdot(\boldsymbol{\sigma}\hat{\mathbf{n}})\big|_\Gamma = f[\psi] = \int ds^\mp\, k(s^\mp - s^\pm)\psi(s^+, s^-) \quad . \tag{9}$$

where $\hat{\mathbf{n}}$ is the outward normal and $\hat{\mathbf{t}}$ is the tangent direction along the shared edge (considered the same direction for both cells).

From dimensional analysis, we can see that:

$$b \sim \frac{1}{t} \quad , \quad D \sim \frac{\ell^2}{t} \quad , \quad \eta \sim F\ell^3 t \quad , \tag{10}$$

where $t$, $\ell$, and $F$ stand for units of time, length, and force. We can then choose times, lengths, and forces to be in units of $\tau$, $\ell_0$, and $F_0$:

$$\tau \equiv \frac{1}{b} \quad , \quad \ell_0 \equiv \sqrt{D\tau} = \sqrt{\frac{D}{b}} \quad , \quad F_0 \equiv \frac{\eta}{\ell_0^3\tau} \quad . \tag{11}$$

Keeping $\ell_0 = \sqrt{D/b}$ as shorthand, we write every parameter, field, and derivative in our system as a dimensionless number (indicated by primes) multiplied by a unit that comprises of $b$, $\ell_0 = \sqrt{D/b}$ and/or $\eta$. We include additionally the side length of a cell, $L_s$, as well as maximum length of linkers $\ell_{\rm M}$ and their typical nucleation range $\ell_{\rm N}$.

$$\begin{aligned}
u =&u'\ell_0 b \quad , \quad \frac{\partial}{\partial x} = \frac{1}{\ell_0}\frac{\partial}{\partial x'} \quad , \quad \frac{\partial}{\partial t} = b\frac{\partial}{\partial t'} \quad , \quad \rho_0 = \rho_0'\frac{1}{\ell_0^2} \quad , \quad \rho = \rho''\rho_0'\frac{1}{\ell_0^2} \\
\gamma =&\gamma'\frac{\eta}{\ell_0^4} \quad , \quad s_1 = s_1'\eta b \quad , \quad s = s'\frac{\eta b}{\ell_0^2} \quad , \quad \nu = \nu'\frac{b}{\ell_0^2} \quad , \quad D \equiv \ell_0^2 b \\
\psi =&\frac{\psi''}{k'}\frac{1}{\ell_0^2} \quad , \quad D_{\rm e} = D_{\rm e}'\ell_0^2 b \quad , \quad s^{\pm} = s^{\pm\prime}\ell_0 \quad , \quad a = a'\frac{b}{\ell_0^2} \quad , \quad \lambda_0 = \lambda_0' b \\
k =&k'\frac{\eta b}{\ell_0^4} \quad , \quad \mu_0 = \mu_0'\frac{\eta}{\ell_0^2} \quad , \quad L_s = L_s'\ell_0 \quad , \quad \ell_{\rm M} = \ell_{\rm M}'\ell_0 \quad , \quad \ell_{\rm N} = \ell_{\rm N}'\ell_0
\end{aligned} \tag{12}$$

Notice that we chose to scale $\rho$ by an additional numerical factor of $\rho_0'$, and we chose to scale $\psi$ by an additional numerical factor of $1/k'$; this is to anticipate algebraic simplifications. Replacing unprimed quantities with primed ones in Eq. (6) gives:

$$\begin{aligned}
\mathbf{0} =&\frac{1}{\ell_0}\nabla'\cdot\left(\eta\frac{1}{\ell_0^4}(\rho''\rho_0'+\rho_0')^2 b(\nabla'\mathbf{u}'+(\nabla'\mathbf{u}')^T) - (s_1'\eta b)\frac{1}{\ell_0^4}(\rho''\rho_0'+\rho_0')^2\mathbb{1} + \left(s'\frac{\eta b}{\ell_0^2}\right)\frac{\rho''\rho_0'}{\ell_0^2}\mathbb{1}\right) \\
&-\left(\gamma'\frac{\eta}{\ell_0^4}\right)\frac{1}{\ell_0^2}(\rho''\rho_0'+\rho_0')\ell_0 b\mathbf{u}' \quad .
\end{aligned} \tag{13}$$

Simplifying gives:

$$\mathbf{0} = \nabla'\cdot\big((\rho''+1)^2(\nabla'\mathbf{u}'+(\nabla'\mathbf{u}')^T) - s_1'(\rho''+1)^2\mathbb{1} + s''\rho''\mathbb{1}\big) - \gamma''(\rho''+1)\mathbf{u}' \quad , \tag{14}$$

where we defined $s'' = s'/\rho_0'$ and $\gamma'' = \gamma'/\rho_0'$. Replacing non-primed quantities with primed ones in Eq. (5), we obtain something similar:

$$b\frac{\partial}{\partial t'}\left(\frac{\rho_0'}{\ell_0^2}\rho''\right) = (\ell_0^2 b)\frac{1}{\ell_0^2}\nabla^{2\prime}\left(\frac{\rho''\rho_0'}{\ell_0^2}\right) + \frac{1}{\ell_0}\nabla'\cdot\left(\ell_0 b\mathbf{u}'\frac{\rho''\rho_0'}{\ell_0^2}\right)) + \nu'\frac{b}{\ell_0^2} - b\frac{\rho_0'}{\ell_0^2}\rho'' \tag{15}$$

in which we substituted $D \equiv \ell_0^2 b$. Eliminating $b\rho_0'/\ell_0^2$ and simplifying, we have:

$$\frac{\partial}{\partial t'}(\rho'') = \nabla^{2\prime}\rho'' + \nabla'\cdot(\mathbf{u}'\rho'') + \nu'' - \rho'' \quad , \tag{16}$$

where we defined $\nu'' = \nu'/\rho_0'$. Continuing to replace non-primed quantities with primed ones in Eq. (8), we can re-write $v^{\pm}$ as:

$$v^{\pm} = (b\ell_0)\left(\frac{k'}{\mu_0'\rho_0'}\frac{(s^{\mp\prime}-s^{\pm\prime})}{(\rho''-1)} + u^{\pm\prime}\right) \equiv (b\ell_0)\left(\frac{1}{\mu_0''}\frac{(s^{\mp\prime}-s^{\pm\prime})}{(\rho''-1)} + u^{\pm\prime}\right) \equiv (b\ell_0)v^{\pm\prime} \quad , \tag{17}$$

where we defined both $v^{\pm\prime}$ and $\mu_0'' \equiv \frac{\mu_0'\rho_0'}{k'}$. We can then replace non-primed quantities with primed ones in Eq. (7) using the above:

$$b\frac{1}{k'\ell_0^2}\frac{\partial\psi''}{\partial t'} = -\frac{1}{\ell_0}\frac{\partial}{\partial s^{+\prime}}\left(-(D_{\rm e}'\ell_0^2 b)\frac{1}{\ell_0}\frac{1}{k'\ell_0^2}\frac{\partial\psi''}{\partial s^{+\prime}} + (b\ell_0)\frac{1}{k'\ell_0^2}v^{+\prime}\psi''\right) - (+\to-) \tag{18}$$

$$+\left(\frac{a'b}{\ell_0^2}\right)\bar{\delta}(|s^{+\prime}-s^{-\prime}|) - (b\lambda_0')\frac{1}{k'\ell_0^2}\psi'' \quad , \tag{19}$$

where we used $(+ \to -)$ as shorthand for the 2nd term of Eq. (7). Eliminating $\frac{b}{k'\ell_0^2}$, we have:

$$\frac{\partial \psi''}{\partial t'} = -\frac{\partial}{\partial s^{+\prime}}\left(-D'_{\mathrm{e}}\frac{\partial \psi''}{\partial s^{+\prime}} + v^{+\prime}\psi''\right) - \frac{\partial}{\partial s^{-\prime}}\left(-D'_{\mathrm{e}}\frac{\partial \psi''}{\partial s^{-\prime}} + v^{-\prime}\psi''\right) + a''\bar{\delta}(|s^{+\prime} - s^{-\prime}|) - \lambda'_0\psi'' , \tag{20}$$

where we defined $a'' = a'k'$. The final equation that requires a non-dimensional form is the boundary condition in Eq. (9). Writing $\boldsymbol{\sigma}$ as a non-dimensional number multiplied by units, we have $\boldsymbol{\sigma} = \boldsymbol{\sigma}'\frac{\eta b}{\ell_0^4}$. Hence Eq. (9) is:

$$\frac{\eta b}{\ell_0^4}\hat{\mathbf{t}} \cdot (\boldsymbol{\sigma}'\hat{\mathbf{n}})\big|_\Gamma = \int ds^{\mp\prime}\ell_0\left(\frac{k'\eta b}{\ell_0^4}\right)\ell_0(s^{\mp\prime} - s^{\pm\prime})\frac{1}{k'\ell_0^2}\psi'' \quad . \tag{21}$$

Simplifying, we have:

$$\hat{\mathbf{t}} \cdot (\boldsymbol{\sigma}^{\pm\prime}\hat{\mathbf{n}})\big|_\Gamma = \int ds^{\mp\prime}\,(s^{\mp\prime} - s^{\pm\prime})\psi'' \tag{22}$$

The elimination of $k'$ and $\rho'_0$ in the final equations is why $\psi$ and $\rho$ were initially scaled with additional numerical factors. Equations (14), (16), and (20) for the fields $\mathbf{u}'$, $\rho''$, and $\psi''$ are the non-dimensional equations that we analyze in this paper. The non-dimensional control parameters required for this model are $s'_1, s'', \gamma'', \nu'', D'_{\mathrm{e}}, a'', \lambda'_0, \mu''_0, L'_s, \ell'_{\mathrm{M}}$, and $\ell'_{\mathrm{N}}$. These are given in terms of the physical, dimensional parameters as the following:

$$\begin{aligned}
s'_1 &= \frac{s_1}{\eta b} \quad , \quad s'' = \frac{s'}{\rho'_0} = \frac{s}{\eta b \rho_0} \quad , \quad \gamma'' = \frac{\gamma'}{\rho'_0} = \frac{\gamma \ell_0^2}{\eta\rho_0} = \frac{\gamma D}{\eta \rho_0 b} \quad , \quad \nu'' = \frac{\nu'}{\rho'_0} = \frac{\nu}{b\rho_0} \\
D'_{\mathrm{e}} &= \frac{D_{\mathrm{e}}}{D} \quad , \quad a'' = a'k' = \frac{ak\ell_0^6}{\eta b^2} = \frac{akD^3}{\eta b^5} \quad , \quad \lambda'_0 = \frac{\lambda_0}{b} \\
\mu''_0 &= \frac{\mu'_0\rho'_0}{k'} = \frac{\mu_0\rho_0 b}{k} \quad , \quad L'_s = \frac{L_s}{\ell_0} = L_s\sqrt{\frac{b}{D}} \quad , \quad \ell'_m = \ell_m\sqrt{\frac{b}{D}} \quad , \quad \ell'_n = \ell_n\sqrt{\frac{b}{D}}
\end{aligned} \tag{23}$$

The non-dimensional boundary conditions are Eq. (22) above along with

$$-\nabla'\rho'' \cdot \hat{\mathbf{n}}\big|_\Gamma = 0 \quad , \quad \mathbf{u}' \cdot \hat{\mathbf{n}}\big|_\Gamma = 0 \quad , \quad -D'_{\mathrm{e}}\frac{\partial \psi''}{\partial s^{\pm\prime}} + v^{\pm\prime}\psi''\big|_{s^{\pm\prime}=0,L'_s} = 0 \quad . \tag{24}$$

From this point on, we will drop primes and double primes for notational clarity. In the main text, all analysis is performed with the non-dimensional form of the equations, hence parameters $D$, $\eta$, $b$, $\rho_0$, and $k$ are all effectively set to 1 whenever not mentioned specifically.

#### C. Choice of parameter values

Since we scaled length on the combination of the diffusion constant $D$ and the time scale $b$ of motors to unbind from F-actin, then a length scale of $\ell_0 = 1$ is interpreted roughly as the length (square root of area) over which F-actin diffuses while it is bound by motors. We expect this length to be small compared to the cell diameter, hence we chose the cell diameter as approximately $16\ell_0$ or $L_s = 8$ for Figs. M1-M3 and $L_s = 16$ for the larger cell in Fig. M4.

We further estimated that dimerized E-cadherin along with their catenins are around 40–80 nm [1] to span between F-actin in one cell and F-actin in the neighboring cell. Cells in the *Drosophila* mesoderm during development are typically around 4–8 $\mu$m in diameter, corresponding to a hexagon side of approximately 2–4 $\mu$m. These measurements imply that the hexagon side should be roughly 25 to 100 times the length of the linker, assuming that the linker is oriented parallel to the membrane. Hence we chose $\ell_\mathrm{M} = 0.6 \approx L_s/13.3$ as the maximum, overstretched linker length and $\ell_\mathrm{N} = 0.1 = L_s/80$ as a typical length for an unstretched linker.

The drag coefficient $\gamma$ of the active gel against an unmodeled substrate is a parameter that when increased significantly, introduces many more complex steady state behaviors to the gel itself, similar to the system in [2]. Hence, in order to focus our analysis on the coupling between cells, we chose to keep $\gamma$ at a low value ($\gamma = 0.001$) so that contractility, steric repulsion, and passive crosslinking are the dominant behaviors in the system, instead of drag against a substrate.

The coefficient of steric force $s_1$ and the contractility strength $s$ were chosen to put the gel in a parameter regime where, with the choice of cell side length $L_s$, contractile dynamics in the system would give rise to condensate formation, that is, distinct regions of high and low density within each cell. An additional consideration is that $\rho$ in the condensate states do not reach extreme numerical levels compared to $\rho_0$.

For linkers, we chose to use a diffusion constant that is low compared to that of F-actin ($D_\mathrm{e} = 0.1$) so that linker movement would be mostly due to being moved by the actin. Additionally, as linkers are embedded in membranes, we expected their diffusion to be lower. The choice of $\mu_0 = 1$ can be roughly interpreted as matching the drag force of linker ends against actin to the spring force from their paired end. We chose this because we did not want one of these forces to dominate the dynamics of the linkers. We ran simulations altering $\mu_0$ (not shown), and have seen some differences in the simulation dynamics, such as arriving at the steady state faster or slower; however, this parameter did not significantly alter the bifurcation diagrams or the emergent coordinated states. The choice of $\lambda_0 = 10 \equiv 10b$ can be interpreted as "fast" linker turnover compared to actin turnover. This is merely a choice, and we ran simulations that varied $\lambda_0$ (not shown) while keeping $a/\lambda_0$ constant. Changing this parameter did alter bifurcation diagrams, but no new globally coordinated states emerged, so this parameter is not emphasized in this work. The nucleation rate $a$ of linkers as a ratio to the degradation rate $\lambda_0$ sets the overall density of linkers. This parameter does alter the bifurcation diagrams, as one can imagine an interpolation between Fig. M**??**(a) and (b) and between Fig. M**??**(a') and (b') (not shown).

### II. TILING GEOMETRIES

#### A. 7-hexagon tile

The minimal repeating unit (polyhex) in which each hexagon has six distinct neighbors consists of seven regular hexagons joined along 21 shared edges. This configuration is technically a *heptahex*, and its periodic, plane-filling arrangement is illustrated in Fig. S1(a).

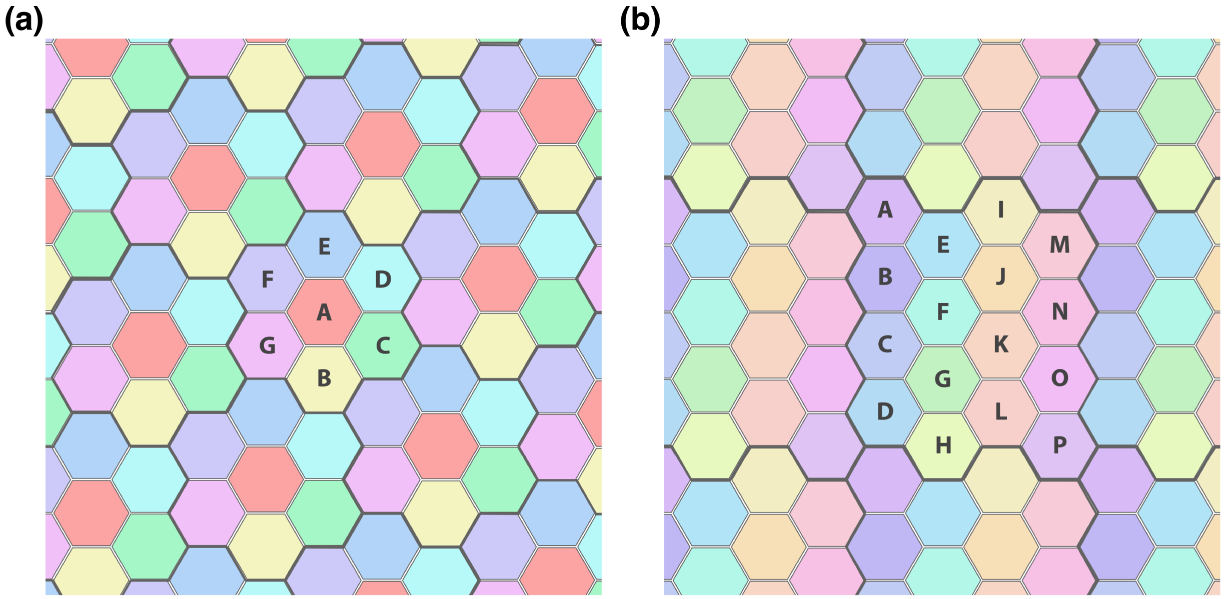


FIG. S1. Schematic of an infinitely periodic system created by repeating a tile of (a) 7 cells (a *heptahex*) and (b) 16 cells across the $xy$ plane.

#### B. 16-hexagon tile

In addition to the odd-numbered heptahex, we consider an even-numbered polyhex consisting of 16 continguous hexagons. This larger tile preserves the property that each cell has six distinct neighbors and its torus-like periodic arrangement is illustrated in Fig. S1(b). As before, each hexagon contributes three distinct edges to the system, yielding a total of 48 associated linker problems.

#### C. Truncated icosahedron: the soccer ball

Our model is general and applies to arbitrary planar geometries; it is not restricted to hexagonal arrays. As an example of a polyhedron with Euler characteristic $\chi = 2$ and mixed polygonal faces, we consider the truncated icosahedron (soccer ball). This polyhedron consists of 32 faces—20 regular hexagons and 12 regular pentagons—joined along 90 edges. Figure S2 depicts the polyhedron with the cell labels and edge-direction conventions used in our simulations. Among the pentagons, six contribute two shared edges to the system (magenta), and six contribute three (yellow), while each hexagon contributes three shared edges (cyan).

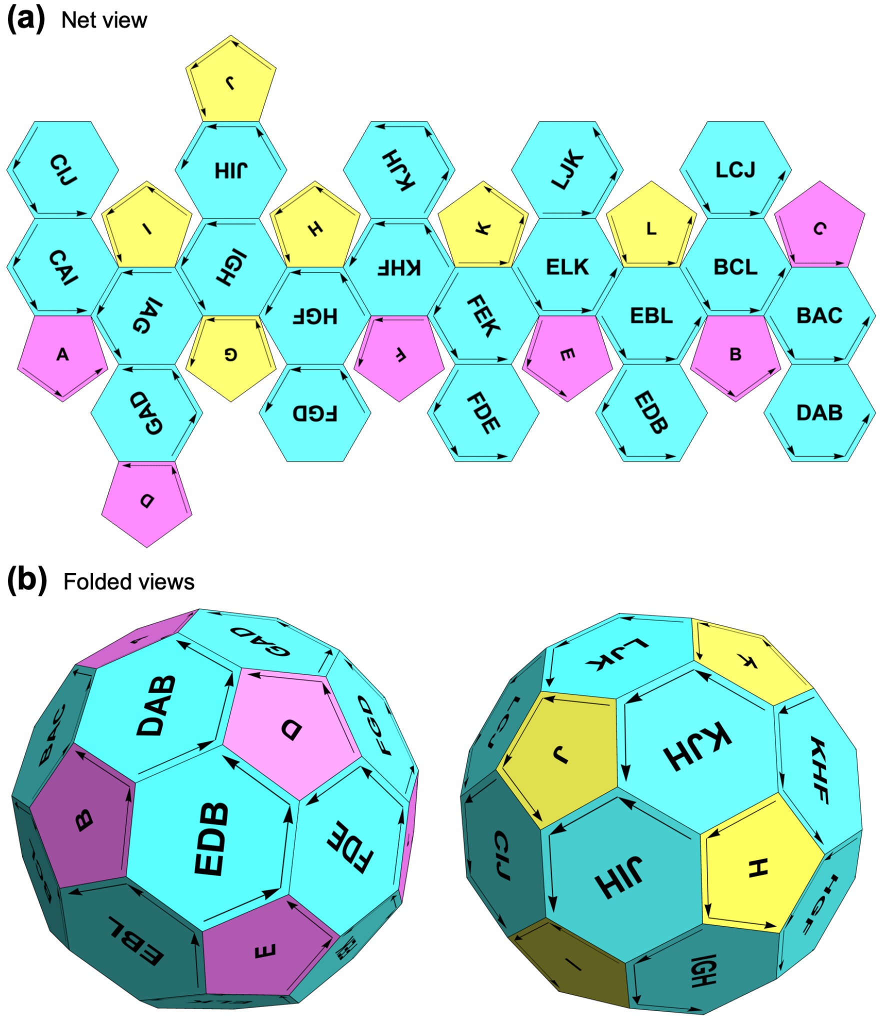


FIG. S2. Schematic of the truncated icosahedron, showing a net view (a) and two folded perspectives (b). The polyhedron consists of (i) six regular pentagons with each contributing two shared edges (magenta, labels A–F), (ii) six pentagons with each contributing three shared edges (yellow, labels G–L), and (iii) 20 regular hexagons with each contributing three shared edges (cyan, labeled according to neighboring pentagons). The arrows indicate the edge-direction convention used in the 90 kinetic problems, shown here on the face of the corresponding "+cell."

### III. COMPUTATIONAL SCHEME

Our system of coupled boundary value problems is solved numerically with a Galerkin finite-element (FE) method using the open-source software FreeFEM++ [3]. We generate two types of computational domains: (i) a polygonal cell domain, as shown in Fig. S3(a), on which each gel is defined, and (ii) a square domain, shown in Fig. S3(b), representing the configuration space of linkers along each shared edge. Both domains are discretized with structured triangular meshes, on which we define FE function spaces based on piecewise linear (P1) basis functions. The evolution of the full coupled system, comprised of gels in cells and linkers on edges, requires transferring and projecting data between the two spaces. To ensure the kinetic problems are properly set up, the orientations of the arc length labels $s^{\pm}$ must share the same convention of positive directionality along each shared edge. We guarantee consistent coupling across the entire system by assigning bottom edges a counterclockwise (CCW) direction and the top edges a clockwise (CW) direction. We note that for tilings of regular hexagons, as in the heptahex or 16-cell tile studied in this paper (Fig. S1), only a single mesh of type (i) is required. For the truncated icosahedron (Fig. S2), we consider a combination of regular hexagons and pentagons with matching edge lengths and uniform edge element densities.

The steps in each time iteration are illustrated in Fig. S3(c) and elaborated below. The system's geometry admits a natural decomposition of the workload, enabling efficient parallel simulation of many cells. In a contiguous polyhex tiling, each hexagon contributes three shared edges to the system. We therefore assign to each CPU process one gel problem in a single cell, together with the linker problems on its three bottom edges. These linker problems exchange data both with the parent cell and with the three neighboring cells below (via MPI). The load distribution for the heptahex tile (Fig. S1(a)) is explicitly outlined in Fig. S3(d) as an example. A similar distribution is applied to the other tilings we consider, with data transfers and projections matching the corresponding cell arrangements. For the truncated icosahedron (Fig. S2), we assign six processes to each handle a pentagon with two edges, six processes to each handle a pentagon with three edges, and 20 processes to each handle a hexagon with three edges (32 cells and 90 shared edges overall).

**Time stepping algorithm**

Hereafter, we omit the primes on dimensionless fields and parameters for clarity. We introduce a time step $dt$, denote the $n^{\text{th}}$ time iteration by superscript, and use the trapezoidal (Crank–Nicolson) rule to integrate the evolution equations for gels and linkers. To implement a Galerkin FE method, we derive the weak (variational) form of each boundary-value

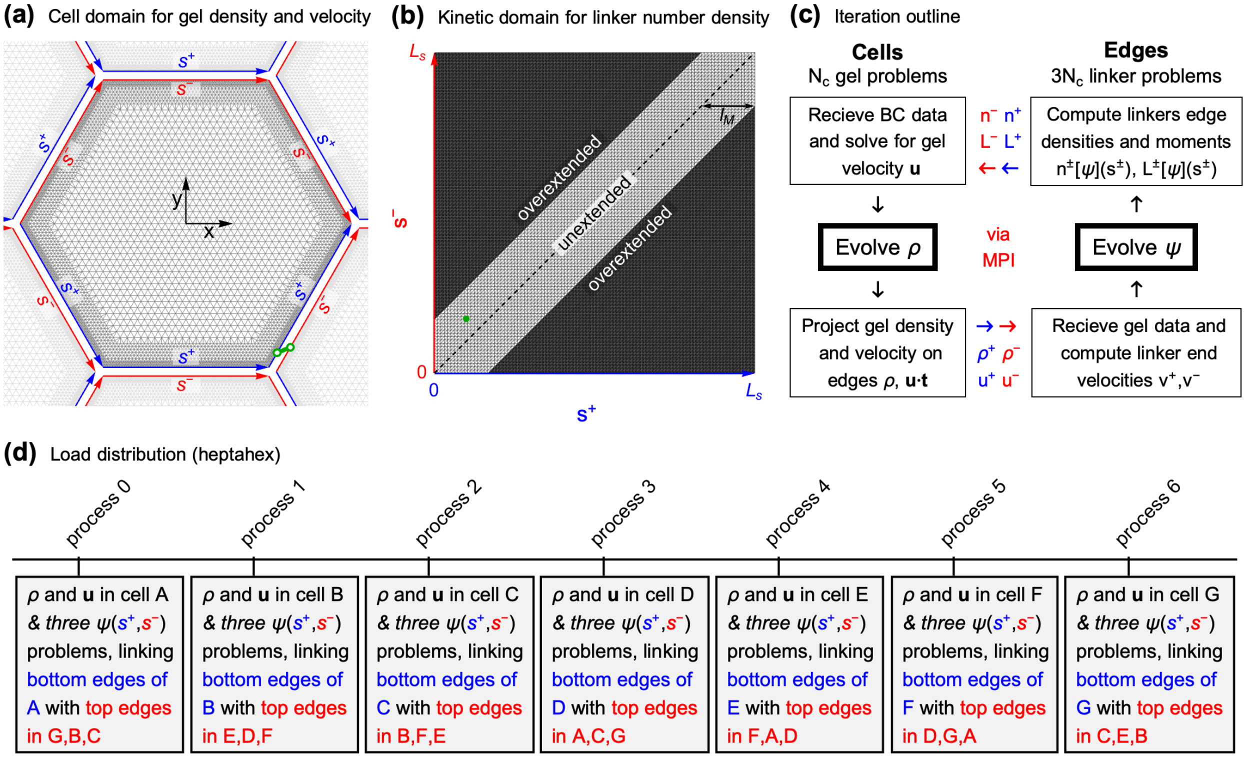


FIG. S3. Computational scheme. (a)–(b) Structured triangular meshes of the hexagonal cell (a) and the linker configuration space (b). Mesh (a) preserves perfect 6-fold symmetry and is refined in layers toward the edges; mesh (b) matches the dimensions and element density of the hexagonal faces in (a). Blue/red arrows indicate arclength labels $s^{\pm}$ with matching directionality along each edge. In (b), the dashed diagonal line marks unextended linker configurations, $|s^+ - s^-| = 0$, and dark regions denote overextended states, $|s^+ - s^-| > l_{\mathrm{M}}$. In green, we illustrate a linker state in physical space (a) mapped to its corresponding point in configuration space (b). (c) For $N_{\mathrm{c}}$ distinct cells, there are as many gel problems to solve on (a) and $3N_{\mathrm{c}}$ linker problems to solve on (b). A time iteration begins with the computation of boundary data to set the flow BCs, and then continues through the steps indicated in the illustrated cycle. Each cell both receives and sends data associated with its six faces. Each kinetic problem defined on (b) exchanges data with the cell above (along $s^+$, blue) and the cell below (along $s^-$, red). (d) Explicit load distribution for the heptahex tile; see Fig. S1(a). The three linker problems associated with the bottom edges of a given cell are handled within the same CPU process, whereas those on the top edges are handled by the processes of neighboring cells.

problem. These are presented below in the order in which they arise in our time-stepping algorithm.

#### 1. Gel momentum balance

To compute the flow $\mathbf{u}$ at time step $n$, we first evaluate the active pressure ($ap$), friction ($fric$), and viscosity ($visc$) using the current gel concentration $\rho^n$.

$$ap^n(\mathbf{x}) = s_1(\rho^n(\mathbf{x})+1)^2 - s\rho^n(\mathbf{x})\,, \tag{25}$$

$$fric^n(\mathbf{x}) = \gamma(\rho^n(\mathbf{x})+1)\,, \tag{26}$$

$$visc^n(\mathbf{x}) = (\rho^n(\mathbf{x})+1)^2\,. \tag{27}$$

Next, we set up the flow BCs by computing appropriate data in the linker configuration domain and transferring it to cell edges. This step must be carried out carefully, as the force signs depend on the chosen convention for edge orientations. In linker configuration space, we compute the following length moments:

$$L^+(s^+) = \int_0^{L_s} ds^-\,(s^+-s^-)\,\psi_i(s^+,s^-)\,, \quad L^-(s^-) = \int_0^{L_s} ds^+\,(s^+-s^-)\,\psi_i(s^+,s^-)\,. \tag{28}$$

These are then transferred to cells and projected along 1D line meshes that outline the polygon edges. The data is carefully transferred to the appropriate edges of appropriate cells, yielding the P1 projections $L_i(\mathbf{x})$ (with $i$ the index of the polygon face and $\mathbf{x}\in\Gamma_i$). Matching Fig. S3(a), the lines at the bottom edges of the hexagon (blue, $i=1,2,3$) are oriented CCW and receive $L^+$ from the corresponding linker problems. The lines at the top of the hexagon (red, $i=4,5,6$) are oriented CW, and receive $L^-$ from the linker problems associated with the neighboring cells above. Following Eq. (M4), and noting that $f^+=-L^+$ and $f^-=L^-$, the boundary condition on the surface force reads:

$$\hat{\mathbf{t}}^{\mathrm{CCW}}\cdot(\boldsymbol{\sigma}\cdot\hat{\mathbf{n}})\Big|_{\Gamma_i} = -L_i \qquad \text{for } i=1,2,3 \text{ (bottom edges)},$$

$$\hat{\mathbf{t}}^{\mathrm{CW}}\cdot(\boldsymbol{\sigma}\cdot\hat{\mathbf{n}})\Big|_{\Gamma_i} = L_i \qquad \text{for } i=4,5,6 \text{ (top edges)},$$

where $\hat{\mathbf{n}}$ is the outward normal. Since $\hat{\mathbf{t}}^{\mathrm{CW}}=-\hat{\mathbf{t}}^{\mathrm{CCW}}$, we have, on all edges,

$$\hat{\mathbf{t}}^{\mathrm{CCW}}\cdot(\boldsymbol{\sigma}\cdot\hat{\mathbf{n}})\Big|_{\Gamma_i} = -L_i\,. \tag{29}$$

To derive the variational form of Eq. (14), we multiply by a vector test function $\mathbf{v}$ (the FE basis functions) and integrate over the cell domain $\Omega$. After integrating the viscous term by parts, we obtain

$$0 = \int_\Gamma \big(visc^n\,(\nabla\mathbf{u}^n + (\nabla\mathbf{u}^n)^T)\cdot\hat{\mathbf{n}}\big)\cdot\mathbf{v} - \int_\Omega visc^n(\nabla\mathbf{u}^n + (\nabla\mathbf{u}^n)^T) : \nabla\mathbf{v}$$
$$- \int_\Omega \nabla ap^n\cdot\mathbf{v} - \int_\Omega fric^n\,\mathbf{u}^n\cdot\mathbf{v}, \tag{30}$$

where $\int_\Gamma$ denotes integration $\int_\Gamma ds$ along the boundary $\Gamma = \partial\Omega$ in the CCW direction, and $\int_\Omega$ denotes the area integral $\iint_{(x,y)\in\Omega} dxdy$. The integrand arising naturally in the first term (the boundary integral) is not determined in full by our boundary conditions. Yet, its tangential component is

$$\hat{\mathbf{t}}^{\mathrm{CCW}}\cdot\big(visc\,(\nabla\mathbf{u} + (\nabla\mathbf{u})^T)\cdot\hat{\mathbf{n}}\big)\Big|_\Gamma = \hat{\mathbf{t}}^{\mathrm{CCW}}\cdot(\boldsymbol{\sigma}\cdot\hat{\mathbf{n}})\Big|_\Gamma,$$

which is specified by Eq. (29). This condition can therefore be imposed in the weak (variational) sense through the tangential projection of the test function along the boundary ($\hat{\mathbf{t}}^{\mathrm{CCW}}\cdot\mathbf{v}$). The normal projection of the test function ($\hat{\mathbf{n}}\cdot\mathbf{v}$) is then used to enforce our additional BC, namely $\mathbf{u}\cdot\hat{\mathbf{n}} = 0$, via penalization. After arranging all bilinear (implicit) terms on the left-hand side [LHS] and all linear (explicit) terms on the right-hand side [RHS], we write the modified variational form as

$$\int_\Gamma M\,(\hat{\mathbf{n}}\cdot\mathbf{u}^n)(\hat{\mathbf{n}}\cdot\mathbf{v}) + \int_\Omega visc^n(\nabla\mathbf{u}^n + (\nabla\mathbf{u}^n)^T) : \nabla\mathbf{v} + \int_\Omega fric^n\,\mathbf{u}^n\cdot\mathbf{v}$$
$$= \int_\Gamma \big(\hat{\mathbf{t}}^{\mathrm{CCW}}\cdot(\boldsymbol{\sigma}\cdot\hat{\mathbf{n}})\big)\,(\hat{\mathbf{t}}^{\mathrm{CCW}}\cdot\mathbf{v}) - \int_\Omega \nabla ap^n\cdot\mathbf{v}, \tag{31}$$

where $M$ is a large numerical parameter penalizing flow penetration, set to $M = 10^6$ in all our simulations. The boundary terms decompose into integrals over the individual faces, all taken with CCW orientation:

$$\int_\Gamma M\,(\hat{\mathbf{n}}\cdot\mathbf{u}^n)(\hat{\mathbf{n}}\cdot\mathbf{v}) = \sum_{i=1}^{6}\int_{\Gamma_i} M\,(\hat{\mathbf{n}}_i\cdot\mathbf{u}^n)(\hat{\mathbf{n}}_i\cdot\mathbf{v}), \tag{32}$$

$$\int_\Gamma \big(\hat{\mathbf{t}}^{\mathrm{CCW}}\cdot(\boldsymbol{\sigma}\cdot\hat{\mathbf{n}})\big)\,(\hat{\mathbf{t}}^{\mathrm{CCW}}\cdot\mathbf{v}) = -\sum_{i=1}^{6}\int_{\Gamma_i} L_i\,(\hat{\mathbf{t}}_i^{\mathrm{CCW}}\cdot\mathbf{v}), \tag{33}$$

where we substituted Eq. (29).

Formally, the variational problem consists of finding $\mathbf{u}^n \in H^1(\Omega)^2$ [the Sobolev space of square-integrable 2D vector fields with square-integrable gradients] such that Eq. (31) holds for all test functions $\mathbf{v} \in H^1(\Omega)^2$. To approximate this problem numerically, we restrict to a finite-dimensional subspace $V_h \subset H^1(\Omega)^2$ spanned by piecewise linear basis functions defined over the triangular mesh $\mathcal{T}_h$ of the cell domain; see Fig. S3(a). This yields a discrete linear system, in which the bilinear form on the LHS of Eq. (31) defines the system matrix. The resulting problem is then inverted using the default *sparsesolver* in FreeFem++.

*2. Gel mass balance*

The variational form of the gel mass balance is obtained by multiplying the time-discretized Eq. (16) by a scalar test function $f$ and integrating over the cell domain. Using the trapezoidal rule and integrating by parts the advection-diffusion terms, we obtain

$$\int_\Omega \Big(\rho^{n+1} f - \frac{dt}{2}\left((\rho^{n+1}\tilde{\mathbf{u}}^{n+1} - \nabla\rho^{n+1})\cdot\nabla f - \rho^{n+1} f\right)\Big) \\ = \int_\Omega \Big(\rho^n f + \frac{dt}{2}\left((\rho^n \mathbf{u}^n - \nabla\rho^n)\cdot\nabla f - \rho^n f\right) + dt\,\nu f\Big), \tag{34}$$

where the boundary terms vanished through substitution of the no-flux condition. For convenience, the LHS collects bilinear terms (defining the matrix) and the RHS collects the linear terms. In the advective term on the LHS, we use $\tilde{\mathbf{u}}^{n+1} = 2\mathbf{u}^n - \mathbf{u}^{n-1}$, a second-order Adams–Bashforth extrapolation of the flow. Within our regime of interest, diffusion was always sufficient to suppress spurious oscillations.

The problem consists of finding $\rho^{n+1} \in H^1(\Omega)$ such that Eq. (34) holds for all test functions $f \in H^1(\Omega)$. As in the previous case, we restrict numerical approximations to the subspace spanned by piecewise linear basis functions defined on the triangular mesh of the cell. We again invert the problem using the default *sparsesolver* in FreeFem++.

*3. Edge projections and linker end velocities*

On the line meshes in the cell space, with directionality convention as in Fig. S3(a), we define the gel's edge density $\rho_i^n(\mathbf{x})$ and edge velocity $u_i^n(\mathbf{x})$ as

$$\rho_i^n(\mathbf{x}) = \rho^n(\mathbf{x})\,, \quad u_i^n(\mathbf{x}) = \begin{cases} \mathbf{u}^n(\mathbf{x})\cdot\hat{\mathbf{t}}^{\mathrm{CCW}} & \text{for } i = 1,2,3 \text{ (bottom edges)} \\ \mathbf{u}^n(\mathbf{x})\cdot\hat{\mathbf{t}}^{\mathrm{CW}} & \text{for } i = 4,5,6 \text{ (top edges)} \end{cases} \,. \quad \text{on } \Gamma_i \tag{35}$$

The data defining these 1D projections are transferred to the appropriate edge problems, where they are interpreted as belonging to either the "+ cell" or the "− cell". In the linker configuration space (Fig. S3(b)), they are defined as $\rho^{\pm,n}(s^{\pm})$ and $u^{\pm,n}(s^{\pm})$.

Similarly, we define the 1D line projections $\rho_i^{n+1}(\mathbf{x})$ and $\tilde{u}_i^{n+1}(\mathbf{x})$ from $\rho^{n+1}(\mathbf{x})$ and $\tilde{\mathbf{u}}^{n+1}(\mathbf{x})$, respectively, to obtain $\rho^{\pm,n+1}(s^{\pm})$ and $\tilde{u}^{\pm,n+1}(s^{\pm})$ in the linker configuration space.

Next, we use these to compute the current linker end velocities:

$$v^{+,n}(s^+,s^-) = \frac{s^- - s^+}{\mu_0\left(\rho^{+,n}(s^+)+1\right)} + u^{+,n}(s^+)\,, \tag{36}$$

$$v^{-,n}(s^+,s^-) = \frac{s^+ - s^-}{\mu_0\left(\rho^{-,n}(s^-)+1\right)} + u^{-,n}(s^-), \tag{37}$$

and their 2nd-order extrapolations to the subsequent time step:

$$\tilde{v}^{+,n+1}(s^+,s^-) = \frac{s^- - s^+}{\mu_0\left(\rho^{+,n+1}(s^+)+1\right)} + \tilde{u}^{+,n+1}(s^+)\,, \tag{38}$$

$$\tilde{v}^{-,n+1}(s^+,s^-) = \frac{s^+ - s^-}{\mu_0\left(\rho^{-,n+1}(s^-)+1\right)} + \tilde{u}^{-,n+1}(s^-). \tag{39}$$

*4. Linkers kinetic problems*

The variational form of the kinetic problem on each shared edge is obtained by multiplying the time-discretized Eq. (20) by a test function $\phi$ and integrating over the linker configuration domain $A$. Using the trapezoidal rule and integrating by parts the advection-diffusion terms, we obtain

$$\begin{aligned}\int_A \Big(&\psi^{n+1}\phi \\ &- \frac{dt}{2}\left((\psi^{n+1}\tilde{v}^{+,n+1} - D_{\mathrm{e}}\partial_{s^+}\psi^{n+1})\partial_{s^+}\phi + (\psi^{n+1}\tilde{v}^{-,n+1} - D_{\mathrm{e}}\partial_{s^-}\psi^{n+1})\partial_{s^-}\phi - k_{\mathrm{off}}\,\psi^{n+1}\phi\right)\Big) \\ &= \int_A \Big(\psi^n\phi + \frac{dt}{2}\left((\psi^n v^{+,n} - D_{\mathrm{e}}\partial_{s^+}\psi^n)\partial_{s^+}\phi + (\psi^n v^{-,n} - D_{\mathrm{e}}\partial_{s^-}\psi^n)\partial_{s^-}\phi - k_{\mathrm{off}}\,\psi^n\phi\right) \\ &\qquad + dt\,k_{\mathrm{on}}\phi\Big),\end{aligned} \tag{40}$$

where $\int_A$ denotes $\int_0^{L_s}\int_0^{L_s} \mathrm{d}s^+\mathrm{d}s^-$ and $k_{\mathrm{off}}(s^+,s^-)$, $k_{\mathrm{on}}(s^+,s^-)$ are *precomputed* P1 interpolations, defined by Eq. (M7). Bilinear forms are collected on the LHS while linear forms are on the RHS. The boundary terms vanished by substituting the no-flux conditions on $s^{\pm}=0$ and $s^{\pm}=L_{\mathrm{s}}$.

The problem consists of finding $\psi^{n+1} \in H^1(A)$ such that Eq. (40) holds for all test functions $\phi \in H^1(A)$. Numerical approximations are restricted to the subspace spanned by piecewise linear basis functions defined on the triangular mesh of the square configuration space (Fig. S3(b)). As before, the LHS defines the matrix and the problem is inverted using the default *sparsesolver* in FreeFem++.

#### 5. *Final updates*

At the end of the time step, we update the following fields:

$$\mathbf{u}^{n-1} = \mathbf{u}^n, \qquad \text{in all cells} \tag{41}$$

$$\rho^n = \rho^{n+1}, \qquad \text{in all cells} \tag{42}$$

$$\psi^n = \psi^{n+1}. \qquad \text{on all edges} \tag{43}$$

This concludes a single time step of our computational algorithm. Example ".edp" scripts for each tile (heptahex, 16-cell, and soccer ball) are provided in the git repository. Output data for $\mathbf{u}$, $\rho$, and $\psi$, together with $n_i$ and $L_i$ (the zeroth and first length moments of $\psi$, interpolated along cell edges), are exported every `nsave` time steps. Since these are all P1 data (amplitudes of "hat function"), the output corresponds to field values at the mesh vertices.

**Post-processing**

We use Mathematica™ for post-processing: importing the various meshes, reinterpolating simulation data onto convenient coordinates, performing quantitative analyses, and generating plots for visualization.

### IV. INCREASED CONTRACTILITY STRENGTH $s$ IN THE ISOLATED CELLS

Increasing the contractility strength $s$ in the isolated cells leads to a split in the polar branch of the steady states. From Fig. S4(a), we see that at the contractility strength $s = 110$, the stable polar branch of the bifurcation diagram contains a break from the values of $\nu \approx 0.22$

to $\nu \approx 0.27$. This is in contrast to Fig. M3(a-a') of the main text where the polar branch is continuous.

From inspection of the density $\rho$ in the polar steady states of the isolated cell, it appears that the now two parts of the polar branch correspond to the gel forming a condensate occupying either one or three corners of the hexagonal domain. Figure S4(b) shows the gel density at the three parameter values indicated with encircled numbers in the bifurcation diagram in (a). The steady state corresponding to parameter set 1 shows the gel condensate occupying one corner of the domain, whereas the steady states corresponding to parameter sets 2 and 3 show the gel condensate occupying three corners of the domain.

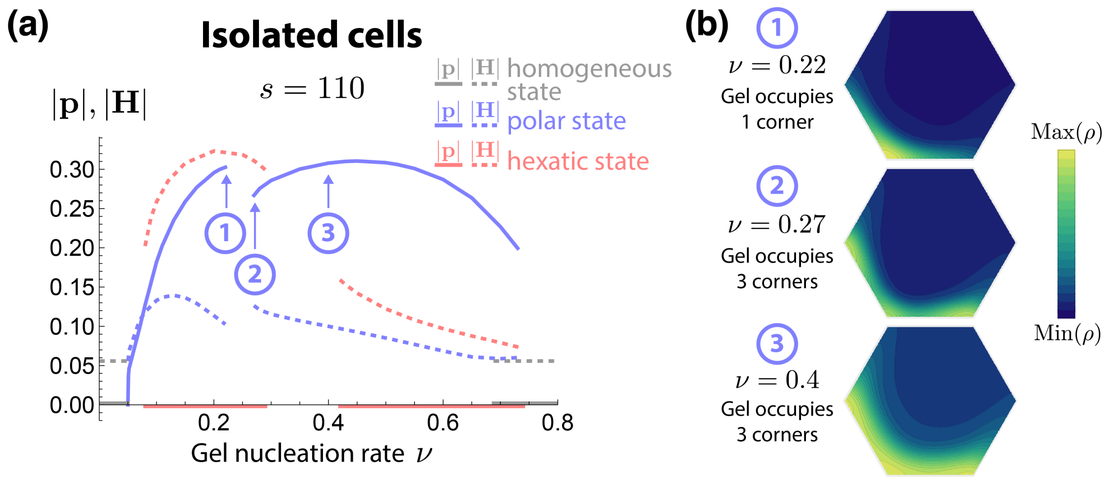


FIG. S4. Higher values of $s$ compared to those in Fig. M3 introduces a split in the polar branch of the steady states for isolated cells. (a) Bifurcation diagram for polar and hexatic states when contractility strength $s = 110$; a split in the polar branch is seen between $\nu \approx 0.22$ and $\nu \approx 0.27$. (b) Gel density profiles $\rho$ at the parameter values indicated by encircled numbers in (a), indicating that the now two parts of the polar branch correspond to the gel occupying either one (parameter set 1) or three (parameter sets 2 and 3) corners of the domain. $s_1 = 30$, $s = 110$, $\gamma = 0.001$. $D = b = \eta = \rho_0 = 1$ from non-dimensionalization.

### V. ABSENCE OF GLOBAL POLARIZATION ON A CLOSED POLYHEDRON (SOCCER BALL)

In polyhex tilings of the plane, we identified a parameter regime in which the fully coupled system robustly converges to a globally polarized state. We then asked what would occur if the same parameters were applied to a system with different geometrical and topological properties. As an illustrative example, we considered the truncated icosahedron (Fig. S2), which has Euler characteristic $\chi = 2$ rather than $\chi = 0$. Such a system cannot support the coordinated polarization of all cells without the formation of defects. The simulation shown in Movie S9, using the same parameters as in Fig. 2(b) and Movie S2, demonstrates this point. Interestingly, the resulting defects are not always sharply defined: while most cells polarize, some instead relax into nematic or hexatic states, and the system ultimately develops an arrested, disordered configuration.

### VI. SUPPLEMENTAL MOVIE LEGENDS

**Movie S1.** Corresponding to Figure 2(a), simulation from random initial conditions showing gel density $\rho$ in 7 isolated cells. One cell evolves into a hexatic steady state; others evolve into polar steady states.

**Movie S2.** Corresponding to Figure 2(b), simulation from random initial conditions showing gel density $\rho$ and linker density $n^{\pm}$ in a 7-cell tile where cells are fully coupled by linkers. The long-term behavior is a globally polarized state steady state.

**Movie S3.** Simulation from random initial conditions with parameters corresponding to Figure 2(c) showing gel density $\rho$; however, the simulation is set in 7 isolated cells instead of linked cells. The hexatic state appears in one out of the 7 cells, but does not spread to other cells.

**Movie S4.** Corresponding to Figure 2(c), simulation from random initial conditions showing gel density $\rho$ and linker density $n^{\pm}$ in a 7-cell tile where cells are fully coupled by linkers. The hexatic state with the actomyosin “ring” spreads across the 7-cell tile.

**Movie S5.** Simulation from random initial conditions showing gel density $\rho$ and linker density $n^{\pm}$ in a 16-cell tile where cells are fully coupled by linkers. Parameters correspond to that of Figure 2(d). The hexatic state with the actomyosin ring visibly spreads across the 16-cell tile, indicating that the global actomyosin ring state is indeed a collective state, the result of cell-to-cell coupling.

**Movie S6.** Corresponding to Figure 2(d), simulation from random initial conditions showing gel density $\rho$ and linker density $n^{\pm}$ in a 7-cell tile where cells are fully coupled by linkers. This movie visually illustrates the tiling the 7-cell tile across the $xy$ plane to make more obvious the nature of the wave-like oscillations shown in the panels of Figure 2(d).

**Movie S7.** Corresponding to Figure 2(f), simulation from random initial conditions showing gel density $\rho$ and linker density $n^{\pm}$ in a 16-cell tile where cells are fully coupled by linkers. This movie shows the evolution of the 16-cell system into a disordered state.

**Movie S8.** Corresponding to Figure 4(a), simulation showing gel density $\rho$ and linker density $n^{\pm}$ in a 7-tile where cells are fully coupled by linkers. This movie shows the dynamics of transient actomyosin cables across the multiple cells. The simulation is started from an initial configuration in which a a gaussian-shaped perturbation is placed at near the center of each cell. At the beginning of the simulation, a star-like pattern of cables appear; this pattern then evolves into multiple cables appearing, disappearing, and moving chaotically throughout the cells.

**Movie S9.** Simulation from random initial conditions showing gel density $\rho$ and linker density $n^{\pm}$ in the truncated icosahedron (soccer ball) system where cells are fully coupled by linkers. Parameters match the heptahex simulation shown in Figure 2(b) and Movie S2. Here, the fully polar state no longer emerges due to the different topology.

**Movie S10.** Corresponding to Figure 4(d), simulation showing gel density $\rho$ and linker density $n^{\pm}$ in the truncated icosahedron (soccer ball) system where cells are fully coupled by linkers. Like Movie S8, this movie shows the dynamics of transient actomyosin cables across a system of multiple confluent cells. At the beginning of the simulation, a star-like pattern of cables appear; this pattern then evolves into multiple cables appearing, disappearing, and moving chaotically throughout the cells.

### REFERENCES

[1] Teiko Shibata-Seki, Masato Nagaoka, Mitsuaki Goto, Eiry Kobatake, and Toshihiro Akaike. Direct visualization of the extracellular binding structure of e-cadherins in liquid. *Scientific Reports*, 10(1):17044, 2020.

[2] Edouard Hannezo, Bo Dong, Pierre Recho, Jean-François Joanny, and Shigeo Hayashi. Cortical instability drives periodic supracellular actin pattern formation in epithelial tubes. *Proceedings of the National Academy of Sciences*, 112(28):8620–8625, 2015.

[3] F. Hecht. New development in freefem++. *J. Numer. Math.*, 20(3-4):251–265, 2012.
